## Supplementary data for "Parallelized manipulation of adherent living cells by magnetic nanoparticles-mediated forces"

### SUPLEMENTARY DATA

**FIGURE S1**

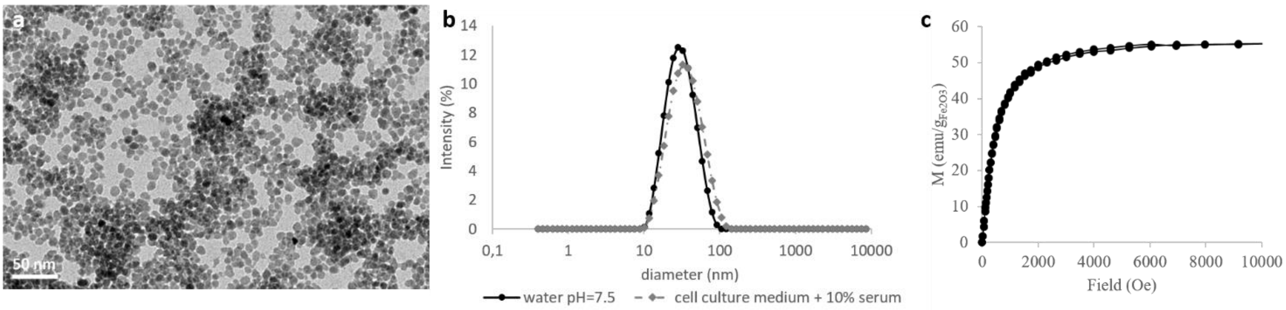


*Figure S1: (a) Transmission electron microscopy image of the $Fe_2O_3$-$PAA_{2k}$-Rhodamine MNPs. The average physical diameter is 8.4 ± 1.9 nm. (b) Dynamic light scattering curves the $Fe_2O_3$-$PAA_{2k}$-Rhodamine MNPs in water (black circles) and in cell culture medium supplemented with 10% serum (grey losange). The mean hydrodynamic sizes of the particles are 27.4 nm and 30.1 in water and in cell culture medium respectively, with a polydispersity index of 0.155 and 0.223. (c) Magnetization curve of the $Fe_2O_3$-$PAA_{2k}$-Rhodamine MNPs. The MNPs are superparamagnetic with a saturation magnetization of 55.7 emu/g, and the magnetic diameter of the particles was measured with a Langevin fit to be 7.96 nm ($\sigma$=0.26).*

**FIGURE S2**

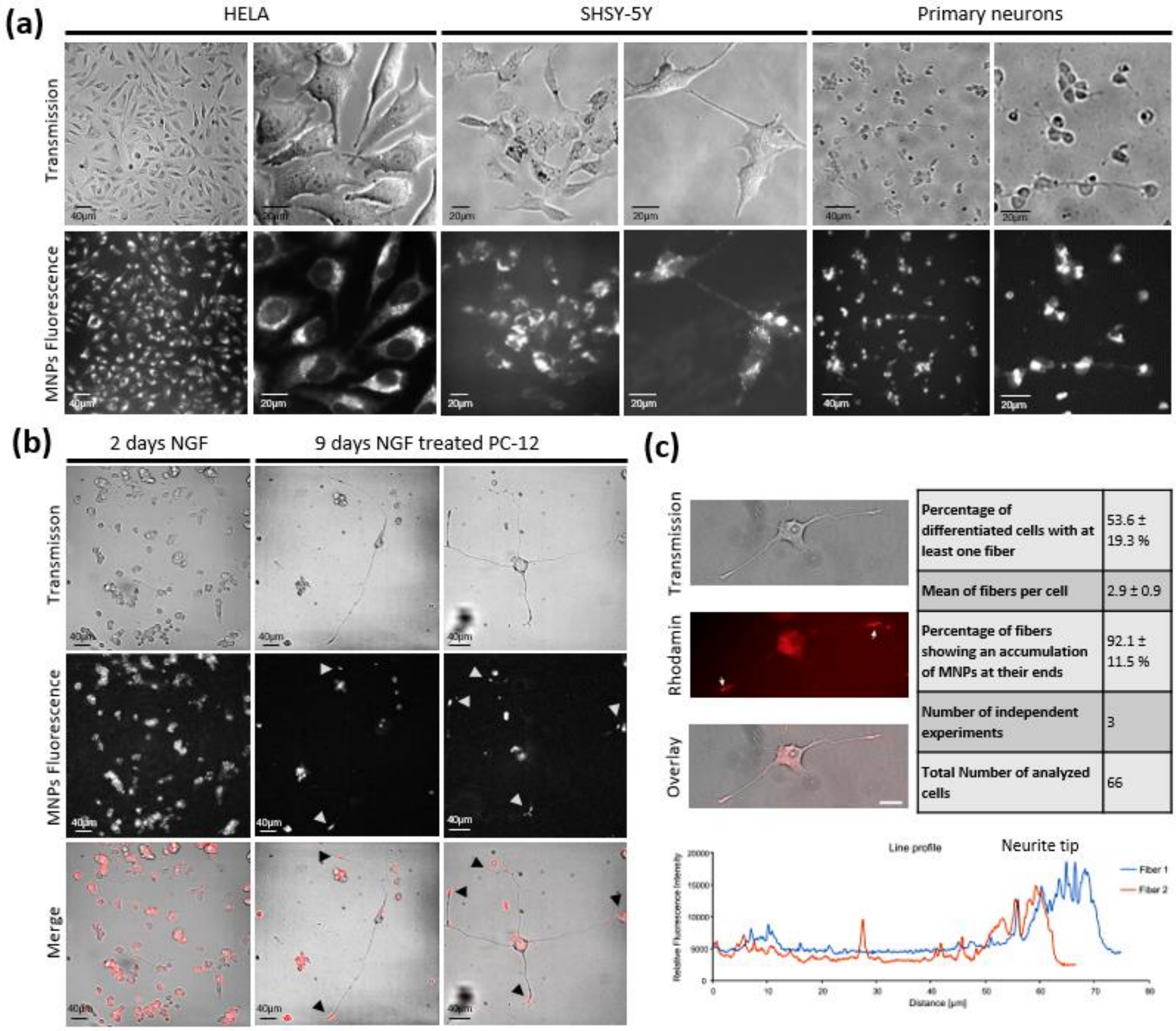


***Figure S2***: *(a) Images of Hela cells, SHSy-5Y cells and primary cortical neurons. The cells were incubated for18-24 hours with* $Fe_2O_3$*-*$PAA_{2k}$*-Rhodamine nanoparticles at a concentration of 10 mM or 2 mM . MNPs were successfully internalized into each cell type. (b) Images of PC12 cells. The cells were incubated 24 hours with MNPs then treated for 2 or 9 days with nerve growth differentiation factor (NGF, 100 ng/ml). As for the other cell type, all PC12 cells have uptake the MNPs. Moreinterestingly MNPs accumulate at the tip of the fibers in a more advanced differentiation state. (c) Amplified images of PC12 cells showing the distribution of MNPs that were taken up for 24 hours and subsequently treated with NGF (100 ng/ml) for 48 hours in differentiation medium in the absence of magnetic forces. 48 hoursTable shows statistical evaluation of MNPs accumulationat the neurite tips. The graph shows the fluorescent intensity (MNPs) along the fibers of a single PC12 cell.*

**FIGURE S3**

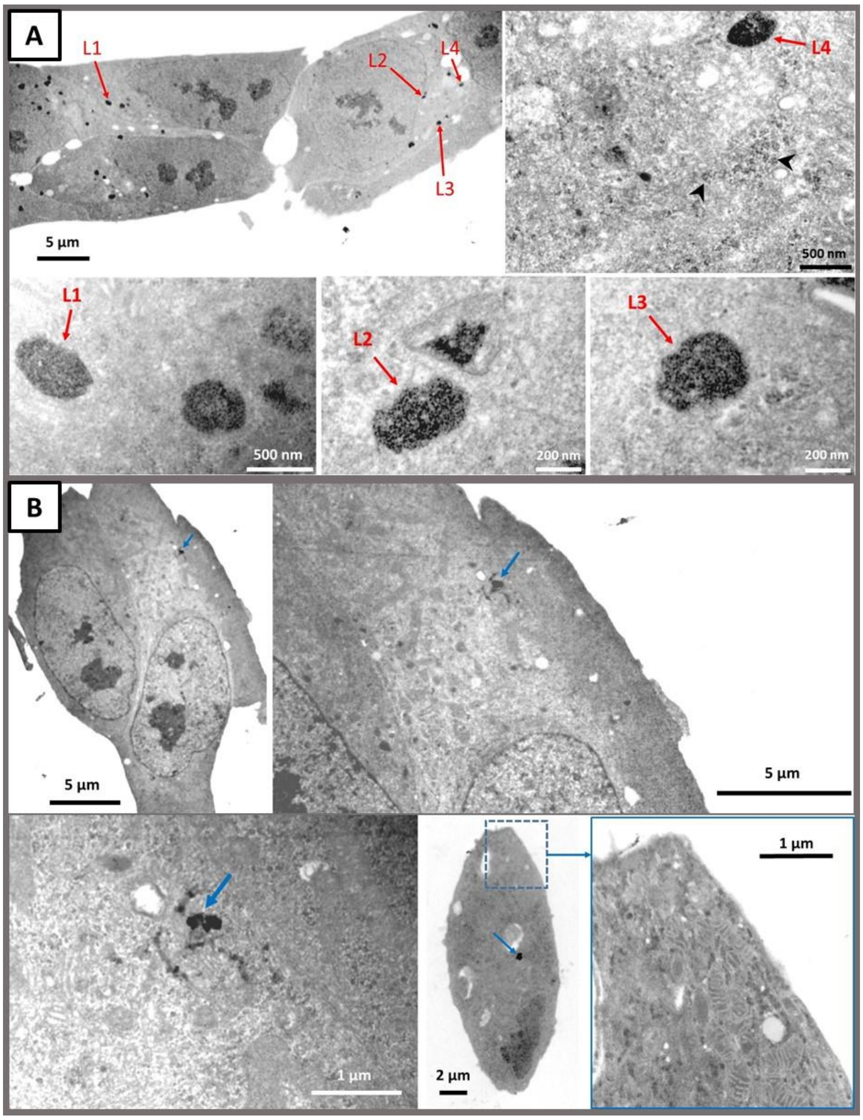


***Figure S3****: Bright field TEM images obtained from 100 nm thick resin embedded sections of SHSy-5Y cells, stained with 2% lead citrate, and 2% uranyl acetate. (A) Cells were incubated with $Fe_2O_3$-$PAA_{2k}$-Rhodamine MNPs for 24 hours. Red arrows indicate MNP-filled lysosomes and black arrowheads show possible MNPs in cytosol. (B) Control sample from cells incubated in the absence of MNPs. Blue arrows show lead citrate deposits.*

| **Cell** | **C1** | **C2** | **C3** | **C4** | **Mean** | **STD** |
|---|---|---|---|---|---|---|
| **Endosome number/section** | 6 | 6 | 10 | 14 | 9 | 3.8 |
| **Endosome number/cell*** | 48 | 48 | 80 | 112 | **72** | **30.6** |

*With section of an endosome ~500 nm and section of SHSy-5Y cell ~4 µm

| Lysosome | L1 | L1’ | L2 | L3 | L4 | Mean | STD |
|---|---|---|---|---|---|---|---|
| **Long diameter , L (nm)** | 602 | 482 | 421 | 392 | 464 | | |
| **Large diameter , l (nm)** | 553 | 371 | 220 | 249 | 272 | | |
| **MNPs number/section** | 260 | 207 | 341 | 408 | 216 | | |
| **100nm endosome volume (nm3)*** | 2,13E+07 | 1,79E+07 | 9,25E+06 | 9,75E+06 | 1,26E+07 | | |
| **MNPs Volumetric density** | 1,22E-05 | 1,15E-05 | 3,68E-05 | 4,18E-05 | 1,70E-05 | | |
| **Endosome volume (nm3)**** | 2,06E+08 | 1,55E+08 | 6,00E+07 | 6,33E+07 | 9,43E+07 | | |
| **MNPs number/endosome** | 2515 | 1790 | 2212 | 2648 | 1611 | **2155** | **448** |

*Volume 100 nm endosome section=L. l. h, with h= 100 nm

**Volume endosome= 4/3. pi. L/2. l/2. ((L/2+l/2)/2). (pi/2)^3

#### FIGURE S4

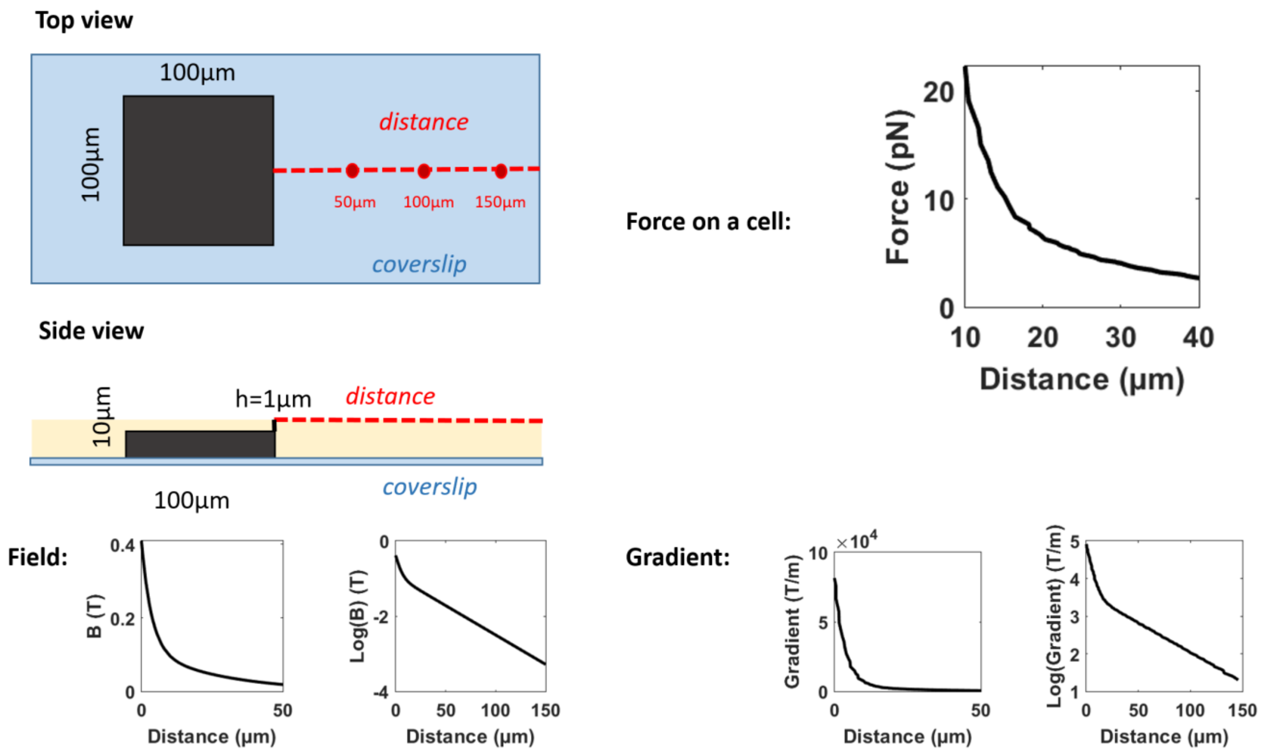


***Figure S4*: Micro-pillar characterization.** *The array is covered with a thin layer of biocompatible polymer (PDMS),slightly thicker than the micropillars, on top of which the cells are platted. The magnetic field (B(T)), the magnetic field gradient (T/m) and the Force per cell (pN) are plotted along a measurement region represented as a red dashed line along the surface of the PDMS layer . The plots were computed using FEM software COMSOL Multiphysics and Matlab. The magnetization curves of both the micropillar and the MNPs were*

*taken into account for those calculations. A typical force of 20 pN is predicted at 10 µm distance from the edge of the micropillar*

**FIGURE S5: MNPs Fate**

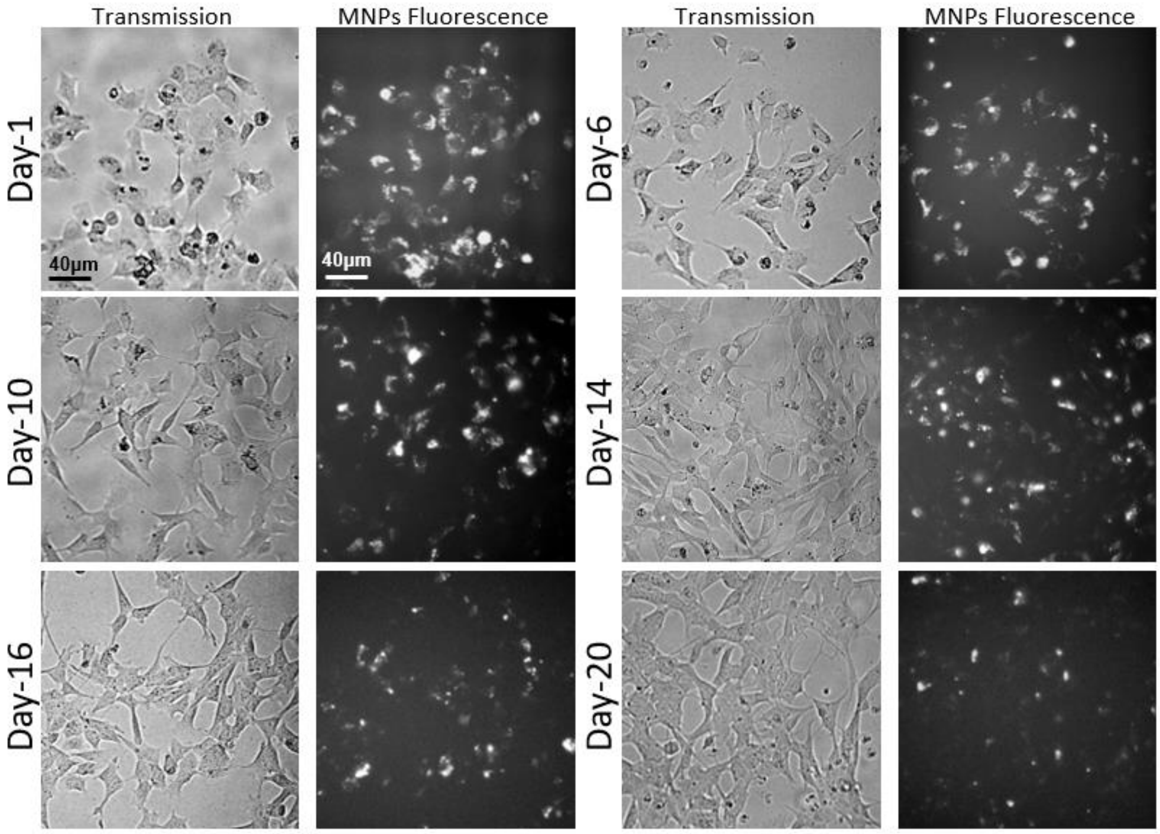


***Figure S5****: Images of SHSy-5Y cells loaded with MNPs. The MNPs were washed away using fresh medium. Cells were then cultured for 20 days under normal culture conditions.*

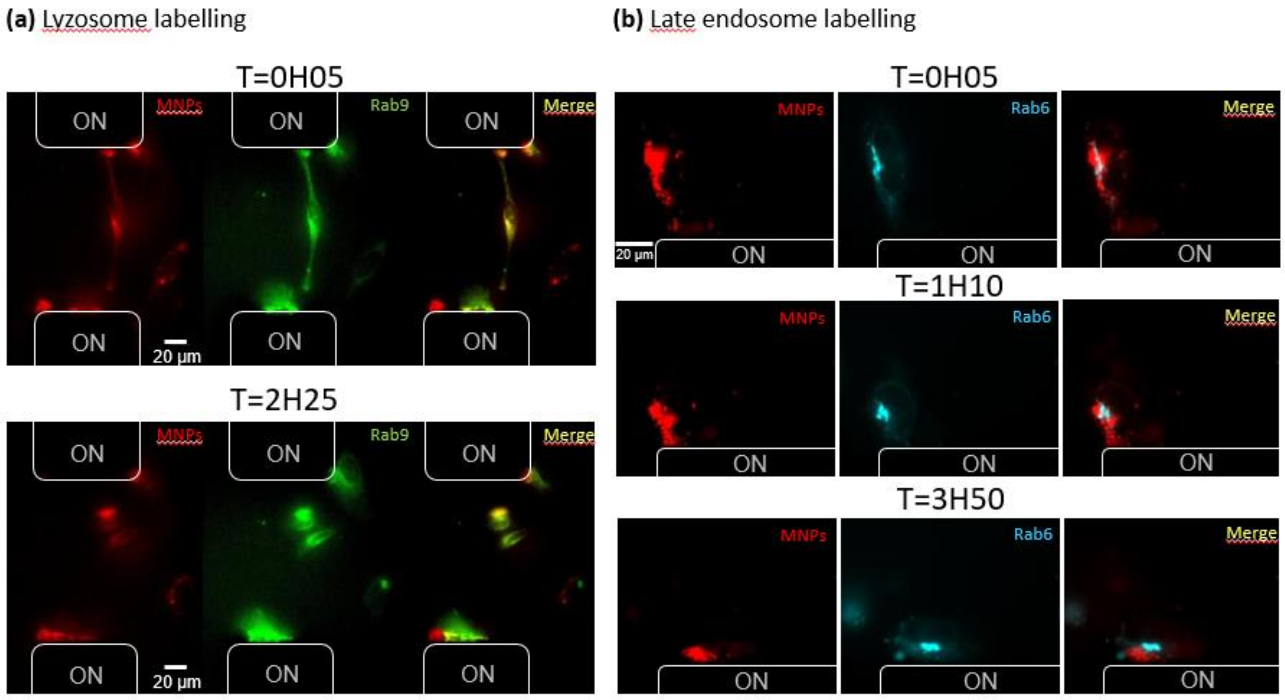

***Figure S5'**: **a) Lysosome labeling**: Images of Hela cells previously transfected with Rab9-GFP (in green) and loaded with $Fe_2O_3$-$PAA_{2k}$-Rhodamine MNPs (in red) on the magnetic array. Pictures show different times of acquisition during 24 hours of magnetic attraction (0H05 and 2H25). Co-localization of particles and lysosomes appears in yellow.*
***b) Late endosome labeling**: Images of Hela cells previously transfected with Rab6-IRFP (in blue) and loaded with $Fe_2O_3PAA_{2k}$-Rhodamine MNPs (in red) on a magnetic array at different times of magnetic attraction (0H05, 1H10 and 3H50).*

**VIDEOS: Voir powerpoint "Supplementary Videos"**

**Video S6: Parallelized magnetic control of Hela cells**

**Video S7: Parallelized magnetic control of SHSY-5Y cells**

**Video S8: Parallelized magnetic control of cortical neurons**

**Video S9: Parallelized magnetic control of cortical neurons**

**Video S10&S11: Parallelized magnetic control of PC12 cells**

**Video S12: Lysosome labeling of Hela cells**

**Video S13: Late endosome labeling of Hela cells**
